# Ultrasound activates mechanosensitive TRAAK K^+^ channels directly through the lipid membrane

**DOI:** 10.1101/2020.10.24.349738

**Authors:** Ben Sorum, Robert A. Rietmeijer, Karthika Gopakumar, Hillel Adesnik, Stephen G. Brohawn

## Abstract

Ultrasound modulates the electrical activity of excitable cells and offers advantages over other neuromodulatory techniques; for example, it can be non-invasively transmitted through skull and focused to deep brain regions. However, the fundamental cellular, molecular, and mechanistic bases of ultrasonic neuromodulation are largely unknown. Here, we demonstrate ultrasound activation of the mechanosensitive K^+^ channel TRAAK with sub-millisecond kinetics to an extent comparable to canonical mechanical activation. Single channel recordings reveal a common basis for ultrasonic and mechanical activation with stimulus-graded destabilization of long-duration closures and promotion of full conductance openings. Ultrasonic energy is transduced to TRAAK directly through the membrane in the absence of other cellular components, likely increasing membrane tension to promote channel opening. We further demonstrate ultrasonic modulation of neuronally expressed TRAAK. These results suggest mechanosensitive channels underlie physiological responses to ultrasound and provides a framework for developing channel-based sonogentic actuators for acoustic neuromodulation of genetically targeted cells.

## Introduction

Manipulating cellular electrical activity is central to basic research and is clinically important for the treatment of neurological disorders including Parkinson’s disease, depression, epilepsy, and schizophrenia^1–4^. Optogenetics, chemogenetics, deep brain stimulation, transcranial electrical stimulation, and transcranial magnetic stimulation are widely utilized neuromodulatory techniques, but each is associated with physical or biological limitations^5^. Transcranial stimulation affords poor spatial resolution; deep brain stimulation and optogenetic manipulation typically require surgical implantation of stimulus delivery systems, and optogenetic and chemogenetic approaches necessitate genetic targeting of light- or small molecule-responsive proteins.

Ultrasound was first recognized to modulate cellular electrical activity almost a century ago and ultrasonic neuromodulation has since been widely reported in the brain, peripheral nervous system, and heart of humans and model organisms^5–12^. Ultrasonic neuromodulation has garnered increased attention for its advantageous physical properties. Ultrasound penetrates deeply through biological tissues and can be focused to sub-mm^3^ volumes without transferring substantial energy to overlaying tissue, so it can be delivered noninvasively, for example, to deep structures in the brain through the skull. Notably, ultrasound generates excitatory and/or inhibitory effects depending on the system under study and stimulus paradigm^5,13,14^.

The mechanisms underlying the effects of ultrasound on excitable cells remain largely unknown^5,13^. Ultrasound can generate a combination of thermal and mechanical effects on targeted tissue^15,16^ in addition to potential off-target effects through the auditory system^17,18^. Thermal and cavitation effects, while productively harnessed to ablate tissue or transiently open the blood-brain barrier^19^, require stimulation of higher power, frequency, and/or duration than typically utilized for neuromodulation^5^. Intramembrane cavitation or compressive and expansive effects on lipid bilayers could generate non-selective currents that alter cellular electrical activity^5,13^. Alternatively, ultrasound could activate mechanosensitive ion channels through the deposition of acoustic radiation force that increases membrane tension or geometrically deforms the lipid bilayer^5,15^. Consistent with this notion, behavioral responses to ultrasound in *C. elegans* require mechanosensitive, but not thermosensitive, ion channels^20^ and a number of mechanosensitive (and force-sensitive, but non-canonically mechanosensitive) ion channels have been implicated in cellular responses to ultrasound including two-pore domain K^+^ channels (K2Ps), Piezo1, Mec-4, TRPA1, TRP-4, MscL, and voltage-gated Na^+^ and Ca^2+^ channels^20–26^. Precisely how ultrasound impacts the activity of these channels is not known.

To better understand mechanisms underlying ultrasonic neuromodulation, we investigated the effects of ultrasound on the mechanosensitive ion channel TRAAK^27,28^. K2P channels including TRAAK are responsible for so called “leak-type” currents because they approximate voltage- and time-independent K^+^-selective holes in the membrane, although more complex gating and regulation of K2P channels is increasingly appreciated^29,30^. TRAAK has a very low open probability in the absence of membrane tension and is robustly activated by force through the lipid bilayer^31–33^. TRAAK activation involves conformational changes that prevent lipids from entering the channel to block K^+^ conduction^32^. Gating conformational changes are associated with shape changes that expand and make the channel more cylindrical in the membrane plane upon opening. These shape changes are energetically favored in the presence of membrane tension, resulting in a tension-dependent energy difference between states that favors channel opening^32^. TRAAK is expressed in neurons and has been localized exclusively to nodes of Ranvier, the excitable action potential propagating regions of myelinated axons^34,35^. TRAAK is found in most (~80%) myelinated nerve fibers in both the central and peripheral nervous system, where it accounts for ~25% of basal nodal K^+^ currents. As in heterologous systems, mechanical stimulation robustly activates nodal TRAAK. TRAAK is functionally important for setting the resting potential and maintaining voltage-gated Na^+^ channel availability for spiking in nodes; loss of TRAAK function impairs high-speed and high-frequency nerve conduction^34,35^. Changes in TRAAK activity therefore appear well poised to widely impact neuronal excitability.

We find that low intensity and short duration ultrasound rapidly and robustly activates TRAAK channels. Activation is observed in patches from cells, from purified channels in reconstituted membranes, and in mouse cortical neurons. Single channel recordings reveal that canonical mechanical- and ultrasonic-activation are accomplished through a shared mechanism. We conclude that ultrasound activates TRAAK through the lipid membrane likely by increasing membrane tension to favor channel opening. This work represents the first demonstration of ultrasound activation of an ion channel directly through the membrane, is consistent with endogenous mechanosensitive channel activity underlying physiological effects of ultrasound, and provides a framework for the development of exogenously expressed sonogenetic tools for ultrasonic control of neural activity.

## Results

We used a recording setup schematized in Figure S1A to isolate mechanical effects of ultrasound mediated through the membrane on ion channels. An ultrasound transducer is connected through tubing to a hole in the recording chamber filled with bath solution. Patched membranes are positioned directly above the transducer face at the position of maximum ultrasonic intensity to eliminate impedance differences between the transducer and membrane and associated surface effects. We designed stimulation protocols to minimize temperature increases (to less than ~O.O5°C, unless otherwise noted).

We first asked whether ultrasound activates TRAAK expressed in cells. As expected, currents from patches pulled from TRAAK-expressing cells were robustly activated by membrane tension created by pressure application through the patch pipette (Figure 1A,B)^28,33^. Strikingly, TRAAK was similarly activated by brief pulses of low intensity ultrasound (10 ms, 5 MHz, 1.2 W/cm^2^). Like basal and pressure-activated TRAAK currents, ultrasound-activated currents were K^+^-selective with a reversal potential near the Nernst equilibrium potential for K^+^ (E_K_^+^ = −59 mV) (Figure 1A,B). Consistent with previous reports, the degree of outward rectification decreased as channel activity increased^36^. Increasing steps of ultrasound power increasingly activated TRAAK current (Figure 1C,D) with a midpoint power of 0.8 ± 0.05 W/cm^2^ and 10%-90% activation occurring between 0.3 and 1.1 W/cm^2^. At the highest ultrasound intensities tested, TRAAK was activated 21.9 ± 5.2-fold (mean ± s.e.m., n=6). Ultrasound did not activate a related, but non-mechanosensitive, K2P ion channel TASK2 (Figure S1C-E).

**Figure 1.**
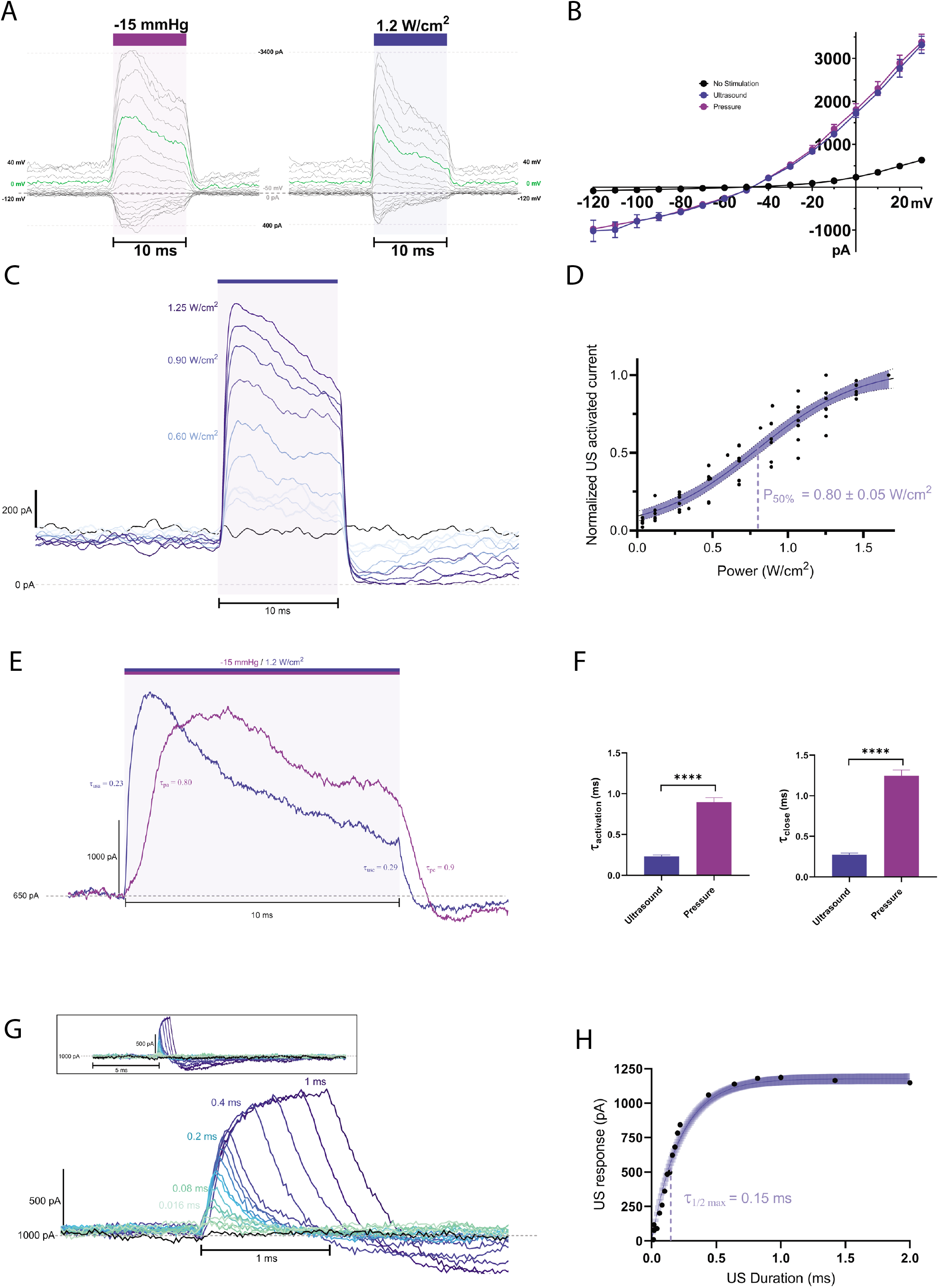
Ultrasound activates TRAAK channels expressed in cell membranes. (A) Currents recorded an excised TRAAK-expressing oocyte patch during a voltage step protocol (V_hold_= −50 mV, V_step_= −120 to +40 mV, ΔV = 10 mV). A pressure (−15 mmHg, purple bar, left) or ultrasound step (1.2 W/cm^2^ at 5 MHz, blue bar, right) was applied during each voltage step. Dashed line indicated zero current and green trace corresponds to V_hold_=0 mV. (B) Current–voltage relationship of data in (A). Average current before stimulation (black) and peak currents during pressure (purple) and ultrasound (blue) stimulation are shown (mean ± SEM, n=3). (C) Overlay of currents during steps of increasing ultrasound power (V_hold_=0 mV). (D) Normalized ultrasound-induced TRAAK current versus ultrasound power (V_hold_=0 mV). Boltzmann fit with 95% CI is shown (n=6). (E) Overlay of TRAAK current from the same patch in response to ultrasound (blue) and pressure (purple) (V_hold_=0 mV). (F) Time constant of channel activation in response to ultrasound (blue) and pressure (purple) (n=5 patches, ****P<0.0001, Welch’s t-test). (G) Overlay of TRAAK response to ultrasonic stimulation of increasing duration colored from green to blue (V_hold_=0 mV). (H) Maximum current response versus stimulus duration (V_hold_=0 mV). Fit with 95% CI is shown.

Ultrasonic and pressure stimulation both result in a rapid increase in current that decays while stimulus is maintained (Figure 1E,F). Similar desensitization of TRAAK following mechanical activation has been described^37^. Ultrasound activates TRAAK currents approximately four times faster than pressure (*τ*_activation, ultrasound_ = 0.23 ± 0.07 ms, *τ*_activation, pressure_ = 0.90 ± 0.06 ms (mean ± s.e.m., n=15)) (Figure 1F). Ultrasound-activated currents similarly decay faster than pressure-activated currents upon stimulus removal (*τ*_close, ultrasound_ =0.27±0.02, *τ*_close, pressure_=1.25±0.07 (n=15). The difference in macroscopic kinetics is at least partially explained by differences in the time required to deliver each stimulus. Pressure increases with a time constant of 1.3 ms in our setup, while an ultrasonic wave with a velocity of ~1500 m/s in solution will arrive at the membrane approximately one hundred times faster. The measured ultrasound activation kinetics therefore more accurately represent intrinsic TRAAK kinetics, while those measured following pressure stimulations are filtered by the pressure clamp device. A consequence of the rapid ultrasonic activation of TRAAK is that even brief stimulation can activate large currents (Figure 1G,H); 0.15 ms and 0.8 ms stimulation result in ~50% and ~95% maximal TRAAK current, respectively.

We note that currents following termination of ultrasound stimulation are typically lower than the basal current for several milliseconds before returning to their baseline level (Figure 1A,E,G) This is similarly observed following pressure stimulation (Figure 1A,E). In the case of pressure, this has been attributed to recruitment of additional lipid into the patch during stretching, which upon stimulus removal results in transiently lower basal tension and reduced channel open probability^33,37^. The similar effect observed following ultrasound and pressure stimulation is consistent with both stimuli generating membrane tension that opens TRAAK channels.

We performed single channel recordings of TRAAK to better understand the basis of channel activation. Consistent with macroscopic records, channel activity was low under basal conditions and increased upon pressure or ultrasound stimulation (Figure 2A,C). Single channels were confirmed to be TRAAK by their K^+^-selectivity, full single channel conductance of ~73 pS at positive potentials, characteristic “flickery” behavior with short openings and multiple subconductance states, and absence in control cells (Figure 2A-C). Ultrasound- and pressure-activated channels opened to an indistinguishable full conductance (Figure 2B). In the absence of stimulation, TRAAK had an average open probability of 1.5±0.9%. Ultrasound and pressure increase channel open probability to 6.2±3.3% and 23±6.8%, respectively. The open probabilities reached upon ultrasound- and pressure-stimulation are not directly comparable because the driving force created by each stimulus was different: high pressures were used to activate channels, but low ultrasound power (0.2 W/cm^2^) was utilized to obtain long (25 s) periods of activation without significant heating. We can place an upper limit on temperature increase during these long stimulations of 0.15 °C. Based on the reported temperature sensitivity of TRAAK (Q10=6), this would result in ~3% activation^31,38^. Thermosensitivity of TRAAK therefore accounts for only a small fraction of the 413 ± 80% increase in open probability upon ultrasound stimulation in these experiments.

**Figure 2.**
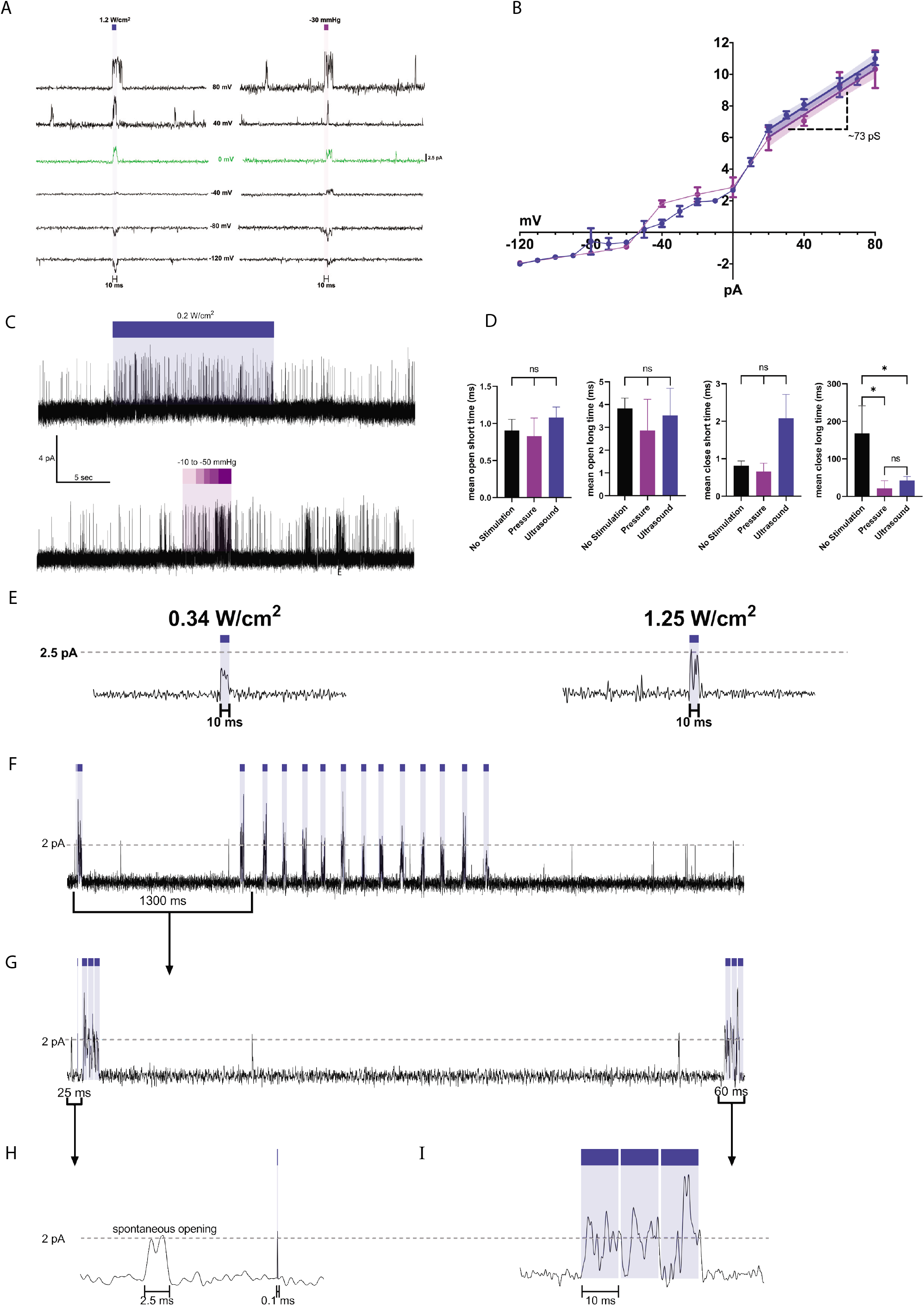
Single TRAAK channel activation by ultrasound. (A) Single TRAAK channel currents recorded at different holding voltages (−120 mV (bottom) to +80 mV (top) ΔV = 10 mV, 40 mV increments shown) with 10 ms stimulation by ultrasound (1.2 W/cm^2^ at 5 MHz, blue bar, left) or pressure (−15 mmHg, purple bar, right). (B) Single channel current elicited by ultrasound (purple) and pressure (blue) versus voltage (n=5 patches). Linear fits from 20 mV to 80 mV with 95% confidence interval and derived conductance is shown. (C) Representative 50 s single channel current with stimulation by ultrasound (0.2 W/cm^2^ at 5 MHz, blue bar, upper) or pressure (−10 to −50 mmHg, purple bar, lower) (V_hold_=0 mV). Long duration ultrasound stimulation was performed at low power to minimize bath temperature increases. (D) Mean open and closed times of TRAAK channels in the absence of stimulation (n=4), during ultrasound stimulation (0.2 W/cm^2^ at 5 MHz, blue, n=6), and during pressure stimulation (~-30 mmHg, purple, n=6). Stimulation significantly decreased mean long closed time (*P < 0.01 for pressure or ultrasound vs. unstimulated, one-way ANOVA) without significantly changing mean open or short closed durations. Mean open (short) times for no stimulation, pressure and ultrasound were 0.90, 0.83, and 1.07 ms. Mean open (long) times for no stimulation, pressure and ultrasound were 3.83, 2.86, and 3.5 ms. Mean closed (short) times for no stimulation, pressure, and ultrasound were 0.81, 0.66, and 2.08 ms. Mean closed (long) times for no stimulation, pressure, and ultrasound were 167.8, 21.4, and 42.5 ms. (E) Single channel current response at 0 mV to steps of increasing ultrasound power (0.34 W/cm^2^ and 1.25 W/cm^2^ at 5 MHz, blue bars, V_hold_=0 mV). Maximum conductance level is indicated with a dashed line. (F-I) Recording demonstrating rapid modulation of channel activity using pulsed ultrasound. (F) A 4.5 s record with multiple periods of ultrasound stimulation (blue bars, V_hold_=0 mV). (G) Magnified view of the 1.3 s portion of the record in (F) indicated with a bar. (H) Magnified view of 25 ms (indicated in (G)) showing a spontaneous and ultrasound-induced opening (100 μs, 0.34 W/cm^2^, 5 MHz). (I) Magnified view of 60 ms (indicated in (G)) showing alternating channel opening and closing in response to pulsed ultrasound (10 ms on, 2.5 ms off, 0.34 W/cm^2^, 5 MHz).

Notably, the major effect of either ultrasonic or pressure stimulation is to increase the frequency of channel opening. TRAAK accesses two open states and two closed states distinguished kinetically (Figure 2D, Figure S3). The mean durations of three states, the two open (with dwell times of ~1 and ~3 ms) and the short duration closed (with a dwell time of ~1 ms), are indistinguishable regardless of whether opening occurs in the absence or presence of stimulation by ultrasound or pressure (Figure 2D). In contrast, the duration of the long-lived closed state is dramatically reduced by either stimulation (from 167.8 ms to 21.4 and 42.5 ms with pressure and ultrasound stimulation, respectively) (Figure 2D).

A subtler effect on channel conductance was also observed upon stimulation (Figure 2E). In addition to a full conductance state, TRAAK frequently opens to subconductance states. Increasing ultrasound power increased the likelihood that a given channel opening would reach full conductance. In the trace shown in Figure 2E recorded at 0 mV, channel openings reached ~1.25 pA (half conductance) with 0.34 W/cm^2^ ultrasound power and ~2.5 pA (full conductance) at 1.25 W/cm^2^. Increasing steps of pressure activation similarly increased the likelihood that openings reached full conductance. Together, these results show pressure and ultrasound open TRAAK channels via a shared mechanism that involves destabilizing long duration closures and favoring full conductance openings with increasing stimulus energy.

We predicted the rapid kinetics of TRAAK activation and ultrasound delivery would permit temporally precise manipulation of single channel activity. In the record shown in Figure 3F, channels activate upon each of 43 distinct bouts of ultrasound stimulation. Closer inspection shows that even very brief (0.1 ms) periods of stimulation activate TRAAK (Figure 2G,H). TRAAK activity was also modulated using pulsed protocols in which longer periods of stimulation (10 ms) are interleaved with brief periods without stimulation (2.5 ms) (Figure 2G,I).

**Figure 3.**
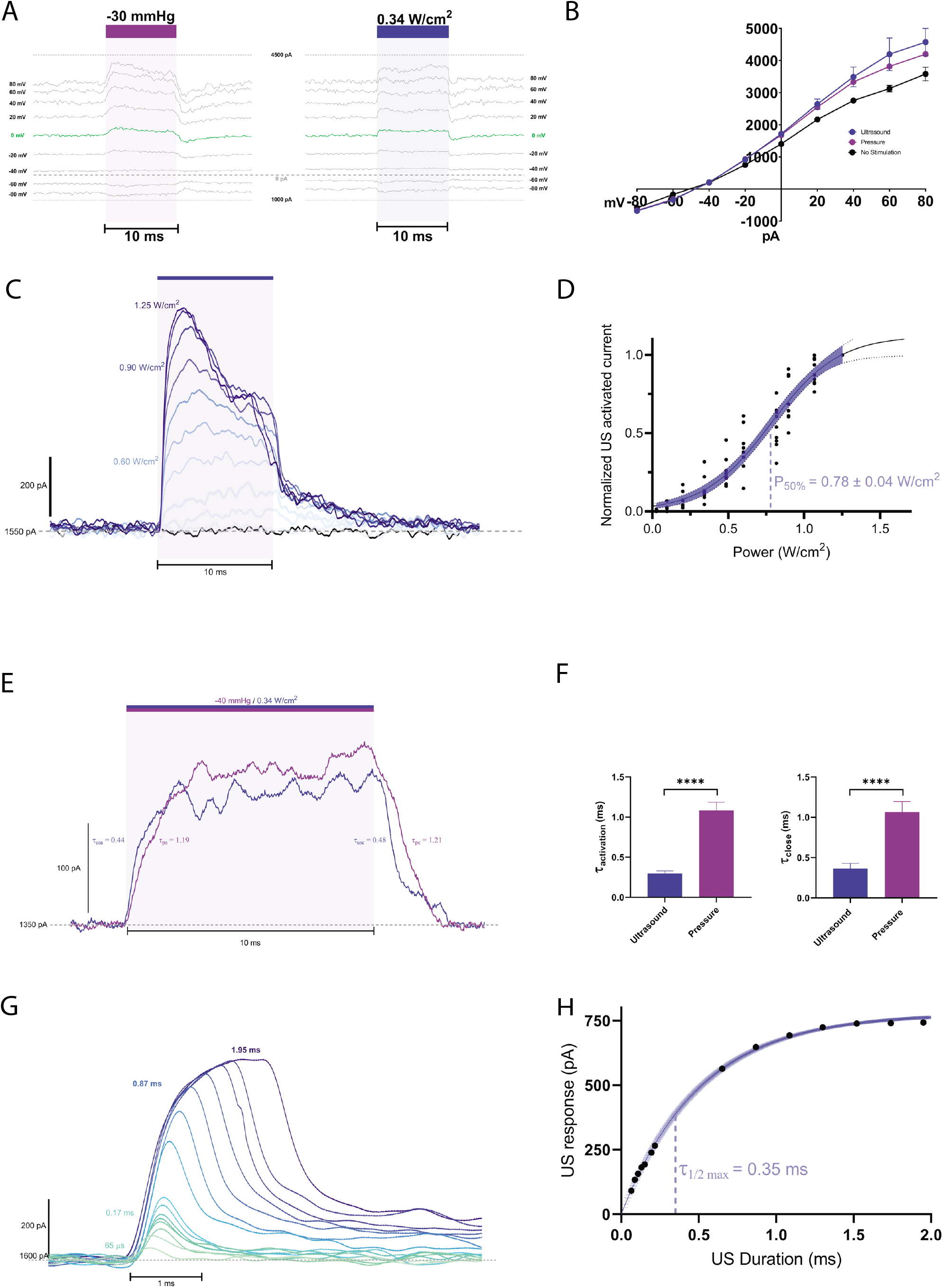
TRAAK channels are activated by ultrasound directly through the membrane. Current recordings from patches of purified TRAAK reconstituted into proteoliposomes. (A) Currents recorded during a voltage step protocol (V_hold_= −50 mV, V_step_= −80 to +80 mV, ΔV = 10 mV, 20 mV increments shown). A pressure (−30 mmHg, purple bar, left) or ultrasound step (0.34 W/cm^2^ at 5 MHz, blue bar, right) was applied during each voltage step. (B) Current–voltage relationship of data in (A). Average current before stimulation (black) and peak currents during pressure (purple) and ultrasound (blue) stimulation are shown (mean ± SEM, n=3). (C) Overlay of currents during steps of increasing ultrasound power colored from light to dark blue (V_hold_=0 mV). (D) Normalized ultrasound-induced TRAAK current versus ultrasound power (V_hold_=0 mV). Boltzmann fit with 95% CI is shown (n=7 patches). (E) Overlay of TRAAK current response from the same patch to ultrasound (blue) and pressure (purple) (V_hold_=0 mV). (F) Time constant of channel activation in response to ultrasound and pressure (n=5 patches, ****P<0.0001, Welch’s t-test). (G) Overlay of TRAAK current response to ultrasound stimulation of increasing duration colored from green to blue (V_hold_=0 mV). (H) Maximum current response versus stimulus duration. Fit with 95% CI is shown.

Ultrasound could in principle activate TRAAK channels in cell membranes in two fundamentally different ways. First, it could activate the channel directly through the lipid membrane by creating membrane tension that favors channel opening. Alternatively, activation could depend on other factors present in cells or on specific components of the lipid membrane. In that case, energy would be conveyed to the channel in a manner analogous to that proposed for mechanosensitive ion channels that require tethers or second messengers^39,40^. To unequivocally distinguish between these possibilities, we studied channel activation in a fully reduced system. TRAAK was heterologously expressed, purified to homogeneity in detergent to remove all other cellular components, and reconstituted into liposomes of defined lipid composition. Proteoliposomes were blistered and high-resistance patches in the inside-out configuration were formed and recorded under voltage clamp. If ultrasound activates TRAAK in this reduced system, it must work through gating forces conveyed to the channel through the membrane.

TRAAK currents from proteoliposomes recapitulated channel properties observed in cellular membranes (Figure 3). Macroscopic currents in the absence of stimulation were K^+^ selective as they reversed near E_K_+ (Figure 3A,B). Pressure steps elicited a rapid increase in current that decayed in the presence of stimulation and rapidly returned to baseline after its removal. As previously reported, the degree of channel activation in this purified system is less than that observed in cells, likely due to an increased tension and basal open probability in reconstituted compared to cellular membranes^33^.

Reconstituted TRAAK was robustly activated by ultrasound (10 ms, 5 MHz, 0.34 W/cm^2^) (Figure 3A,B). Like basal and pressure-activated currents, ultrasound-activated currents were K^+^-selective. Increasing steps of ultrasound power resulted in progressive activation of TRAAK, with a midpoint of ultrasound power activation at 0.78 ± 0.04 W/cm^2^ and 10%-90% activation observed between 0.2 and 1.25 W/cm^2^ (Figure 3C,D). Ultrasound activation and subsequent channel closure proceeded at faster rates than those observed with pressure (Figure 3E,F, *τ*_activation, ultrasound_ = 0.30 ± 0.03 ms, *τ*_activation, pressure_ = 1.08 ± 0.06 ms, *τ*_close, ultrasound_ = 0.37 ± 0.06 ms, *τ*_close, pressure_ = 1.07 ± 0.13 ms (mean ± s.e.m., n=15-33)). As in cells, a consequence of rapid ultrasound activation is that even brief stimulation can effectively activate channels: ~50% and ~95% maximal recruitment is achieved with 0.35 ms and 1.30 ms stimulation, respectively (Figure 3G,H). Recordings from cell membranes and from proteoliposomes show similar power responses and channel kinetics (Figure 1D,F and 3D,F), suggesting the same process underlies channel activation in cells and in purified systems. We conclude that ultrasound activation of TRAAK does not require additional cellular components as sensors or conveyors of energy to the channel. Ultrasound activates TRAAK directly through the lipid membrane.

We then asked whether TRAAK expressed in neurons could be activated by ultrasound. Mice were in utero electroporated with a TRAAK-encoding plasmid and brain slices were harvested from juvenile animals for recording. Confocal images of brain slices showed expression and membrane localization of TRAAK channels in cortical layer 2/3 pyramidal neurons (Figure 4A). Neurons were patched in the whole-cell configuration and recorded in voltage clamp.

**Figure 4.**
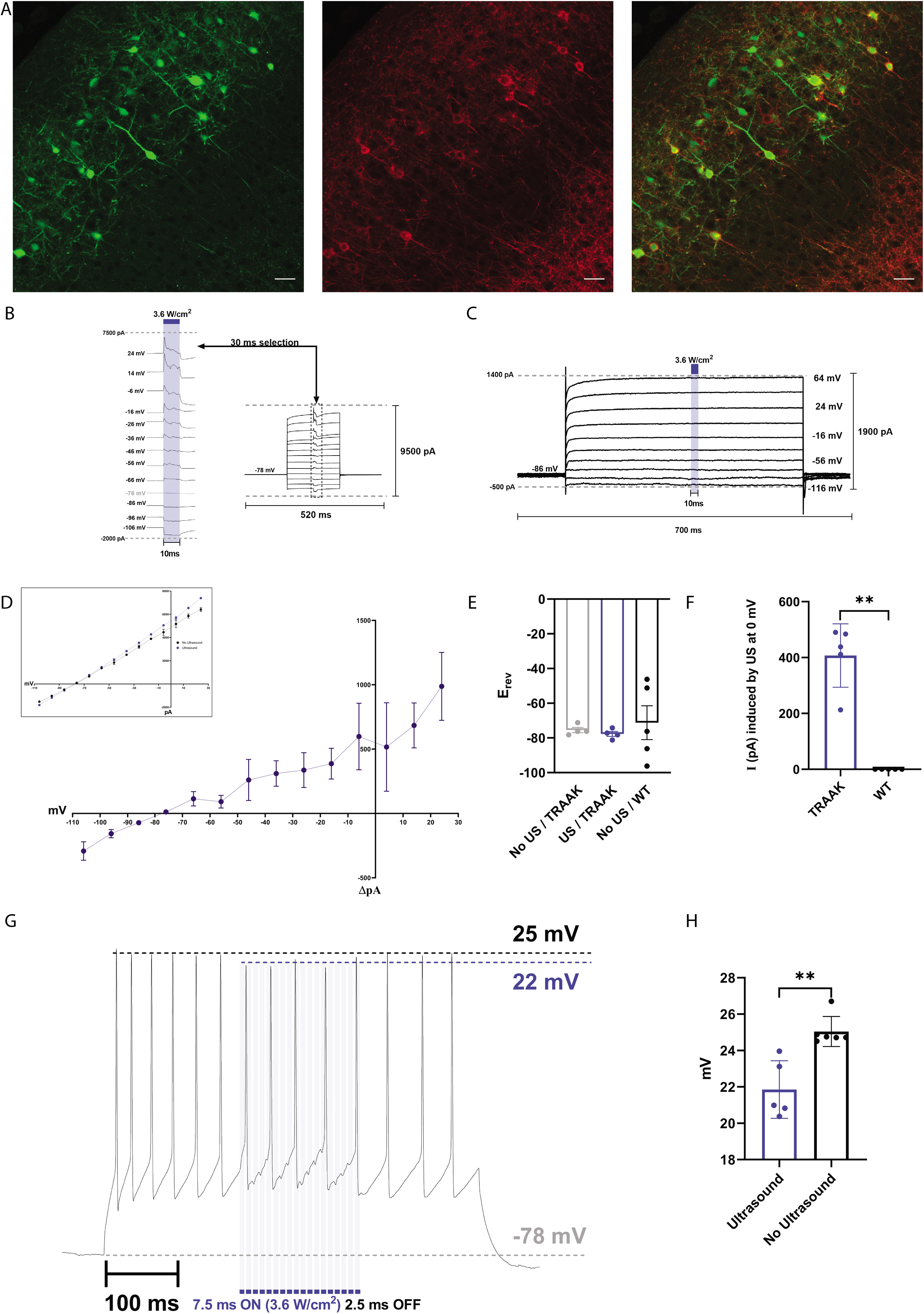
Ultrasound activation of TRAAK channels expressed in mouse brain. (A) Representative confocal images a juvenile mouse cortex co-in utero electroporated with soluble GFP (green, left) and membrane localized TRAAK-mRuby2 (red, center). Merged image is shown at right. (B) Representative whole cell current recordings from a cortical layer 2/3 pyramidal neuron expressing TRAAK during a voltage step protocol (V_hold_ = −78 mV, V_step_= −108 to +24 mV). An ultrasound step (3.6 W/cm^2^ at 5 MHz, 10 ms, blue bar) was applied during each voltage step. (C) Representative control whole cell current recordings from a non-TRAAK expressing cortical layer 2/3 pyramidal neuron during a voltage step protocol (V_hold_ = −86 mV, V_step_= −116 to +64 mV). An ultrasound step (3.6 W/cm^2^ at 5 MHz, 10 ms, blue bar) was applied during each voltage step. (D) Current voltage relationship of ultrasound-activated currents from TRAAK-expressing neurons (mean ± SEM, n=3). Average current before and peak current during ultrasound stimulation is plotted in the inset. (E) Reversal potential of currents recorded from control and TRAAK-expressing neurons in the presence and absence of ultrasound stimulation (n=5, mean ± s.e.m.). (F) Peak ultrasound-activated current at 0 mV recorded from control and TRAAK-expressing neurons (n=5, mean ± s.e.m., **P<0.01). (G) Spike train elicited by current injection (125 pA) from a TRAAK-expressing neuron. Pulsed ultrasound stimulation (7.5 ms on (blue bars), 2.5 ms off, 3.6 W/cm^2^) was applied during the current injection. (H) Mean spike amplitude in the presence and absence of ultrasound stimulation (n=6, mean ± s.e.m., **P<0.01).

Ultrasound stimulation (10 ms, 5MHz, 3.2 W/cm^2^) activated large currents in TRAAK-expressing (Figure 4B,D), but not in control (Figure 4C), cells with a mean activation of ~400 pA at 0 mV (Figure 4F). Ultrasound-stimulated currents in TRAAK expressing cells had rapid opening and closing kinetics (Figure 4B) and negative reversal near E_K_+ (Figure 4D,E), consistent with activation of exogenously expressed TRAAK channels. During a spike train elicited by current injection, pulsed ultrasonic stimulation resulted in phase-matched hyperpolarization during interspike intervals and a ~3 mV reduction in spike amplitude in the presence of stimulation (Figure 4G,H). We note higher power ultrasound was required to activate channels in whole-cell recordings from neurons in brain slice compared to excised patches from cells or proteoliposomes. This may be due to differences in recording setup (Figure S1B), patch-configuration, or the membrane environment in which TRAAK is embedded. These results demonstrate ultrasound can be used to manipulate the activity of TRAAK channels in neurons in the brain.

## Discussion

Here we demonstrate that ultrasound activates mechanosensitive TRAAK ion channels directly through the lipid membrane. Ion channels have been increasingly implicated in mediating the cellular electrical effects of ultrasound in excitable cells. Using microbubbles to amplify the acoustic radiation forces enabled ultrasound activation of the mechanosensitive channels *M. musculus* Piezo1, *E. coli* MscL, and *C. elegans* TRP-4 in cells^25,41,42^. Piezo1 and MscL can be activated by ultrasound in the absence of microbubbles using higher frequency and power or higher frequency and a sensitizing mutation, respectively^22,26^. Behavioral responses of *C. elegans* to ultrasound require the mechanosensitive Mec-4 channel^20^. Non-canonically mechanosensitive channels have also been implicated in cellular responses to ultrasound. Astrocytic TRPA1 accounts for some behavioral and cellular responses to low frequency, low intensity ultrasound (0.4 MHz, 0.3 W/cm^2^) in mice^23^. Voltage-gated channels, some of which are mechanically sensitive, have been implicated in neural responses to ultrasound^24,43^, although other studies report modest effects that may be explained by ultrasound-induced temperature changes^21,22^. Other tension-gated mechanosensitive channels could be expected to be similarly ultrasound-sensitive^39^.

These results here contrast with a previous report of ultrasound activation of TRAAK and related TREK channels in oocytes^21^. In the current study, brief duration and low power stimulation (1.2 W/cm^2^, 5 MHz, 10 ms) robustly activates TRAAK (up to ~20 fold) with fast (*τ* ~ 250 us) kinetics. In previous work, long duration and higher power stimulation (2 W/cm^2^, 5 MHz, 1s) modestly activated channels (up to ~0.15 fold) with thousand-fold slower kinetics (*τ* ~ 800 ms)^21^. Ultrasound activation of TRAAK reported here closely corresponds to canonical mechanical activation in whole cells, patches from cells or proteoliposomes, and single channel recordings.

What is the physical basis for ultrasound activation of TRAAK? We can exclude temperature increase, cavitation, displacement, and acoustic scattering for the following reasons. The stimulation protocols used here result in minimal heating, consistent with previous reports^20,21,23,25^. Cavitation requires approximately ten-fold higher pressures and would open non-selective holes in the membrane, rather than K^+^-selective paths. The expected displacement gradient is small (~0.1 um) over the ~300 um ultrasound wavelength. Similarly, scattering by the glass pipettes likely does not substantially change the ultrasound intensity profile since the tip diameter is small (1 um) relative to ultrasound wavelength. As has been postulated^5,12,22^, acoustic radiation forces and resulting acoustic streaming most likely result in channel activation. We suspect that energy from ultrasound ultimately increases membrane tension to open TRAAK channels. Future efforts to directly measure tension during stimulation could provide insight into protocols that maximally increase tension and optimally activate channels.

Ultrasound has both suppressive and stimulatory effects on neuronal activity, depending on the stimulus design and tissue under study. Inhibitory effects of ultrasound have been demonstrated in the central and peripheral nervous systems including in studies of light evoked potentials in visual cortex, pupillary reflexes, spreading cortical depression, and sciatic nerve. The underlying molecular mechanisms for these effects are unknown. Our results suggest that TRAAK and TREK1 mechanosensitive K^+^ channels are responsible for some ultrasound-induced inhibition of neuronal activity. TRAAK and TREK1 channels are localized to nodes of Ranvier within myelinated axons and their activation is expected to impact spiking by increasing resting K^+^ conductance and hyperpolarizing cells^34,35^. Focused ultrasound stimulation of myelinated fibers containing TRAAK and TREK may be a viable strategy for targeted suppression of neural activity.

An alternative to manipulating endogenously expressed channels is to sensitize targeted cells with overexpression of an ultrasound-activated protein. Several such “sonogenetic” approaches have been reported using the ion channels TRP-4, MscL, and Piezo as ultrasound actuators for neuromodulation or expression of reporter genes^25,26,41,42,44^. Our results provide a framework for the development of TRAAK or other mechanosensitive ion channels as modular tools for targeted suppression or activation of electrically excitable neurons or other cells. The low resting open probability and relatively high conductance (compared to channelrhodopsins used for optogenetics^45^) of TRAAK make it a promising target for further engineering to optimize overexpression, subcellular localization, and ultrasound-responsive range for sonogenetic applications.

---

**Figure S1.**
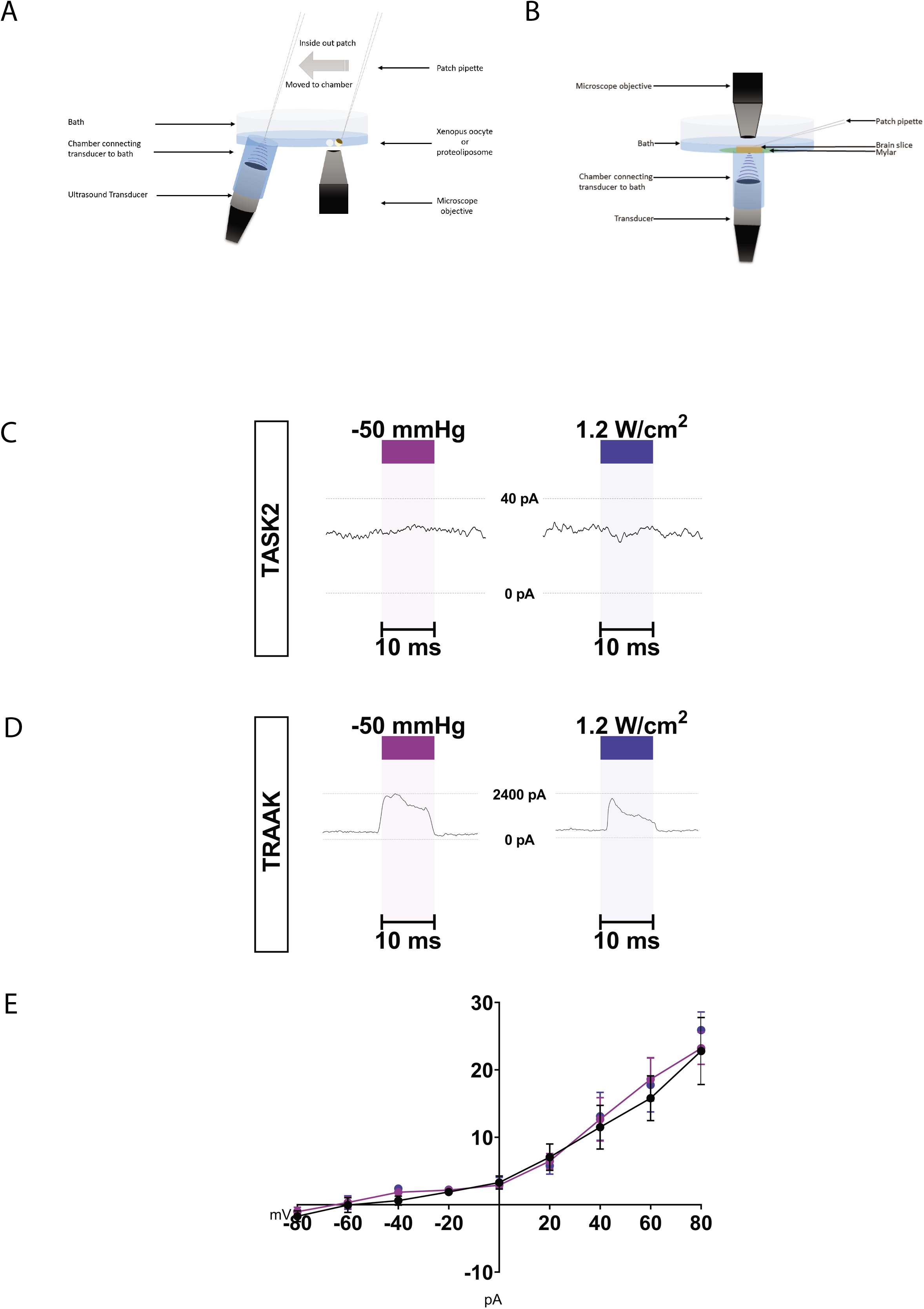
Setup for recording ultrasound effects on ion channels. (A) Chamber design for patch recording experiments. An ultrasound transducer is connected to the bath solution through tubing such that an unobstructed liquid column exists between the transducer face and patch pipette. (B) Chamber design for slice recording experiments. An ultrasound transducer is connected to the bath solution through tubing and a thin mylar sheet on which the brain slice is fixed. (C) Voltage-clamp recording of the non-mechanosensitive K2P channel TASK2 (V_hold_=0 mV). Neither pressure or ultrasound activates TASK2 currents. (D) Voltage-clamp recording of the mechanosensitive K2P channel TRAAK (V_hold_=0 mV). Pressure and ultrasound activate TRAAK currents. (E) Current-voltage relationship from TASK2-containing patches. Average current in the absence of stimulation (black) and maximum currents during pressure (purple) and ultrasound (blue) stimulation are shown (mean ± SEM, n=3). Recordings were made in a 10-fold [K^+^] gradient and presented in physiological convention: E_K+_=-59 mV and positive current indicates K^+^ flow from the high [K^+^] (intracellular) to low [K^+^] (extracellular) side.

**Figure S2.**
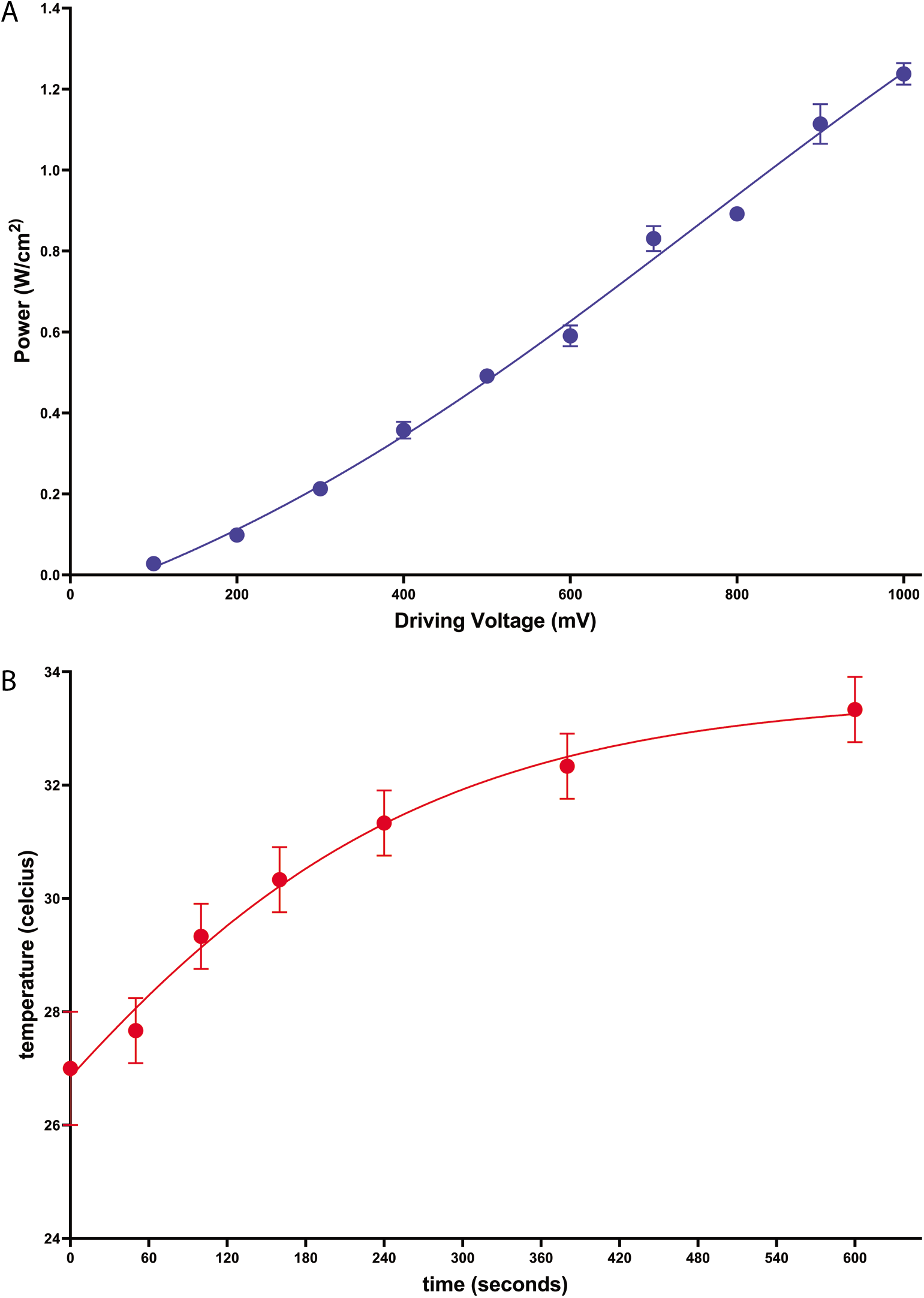
Calibration of ultrasound power and temperature increases. (A) Power versus driving voltage. Pressure was measured at the point of maximum ultrasound intensity using a manufacturer-calibrated needle hydrophone and converted to power as described in Methods. (B) Temperature increase versus ultrasound stimulation time. Temperature increases at the point of maximum ultrasound intensity were made using a thermocouple during constant stimulation at the maximum driving voltage used as described in Methods.

**Figure S3.**
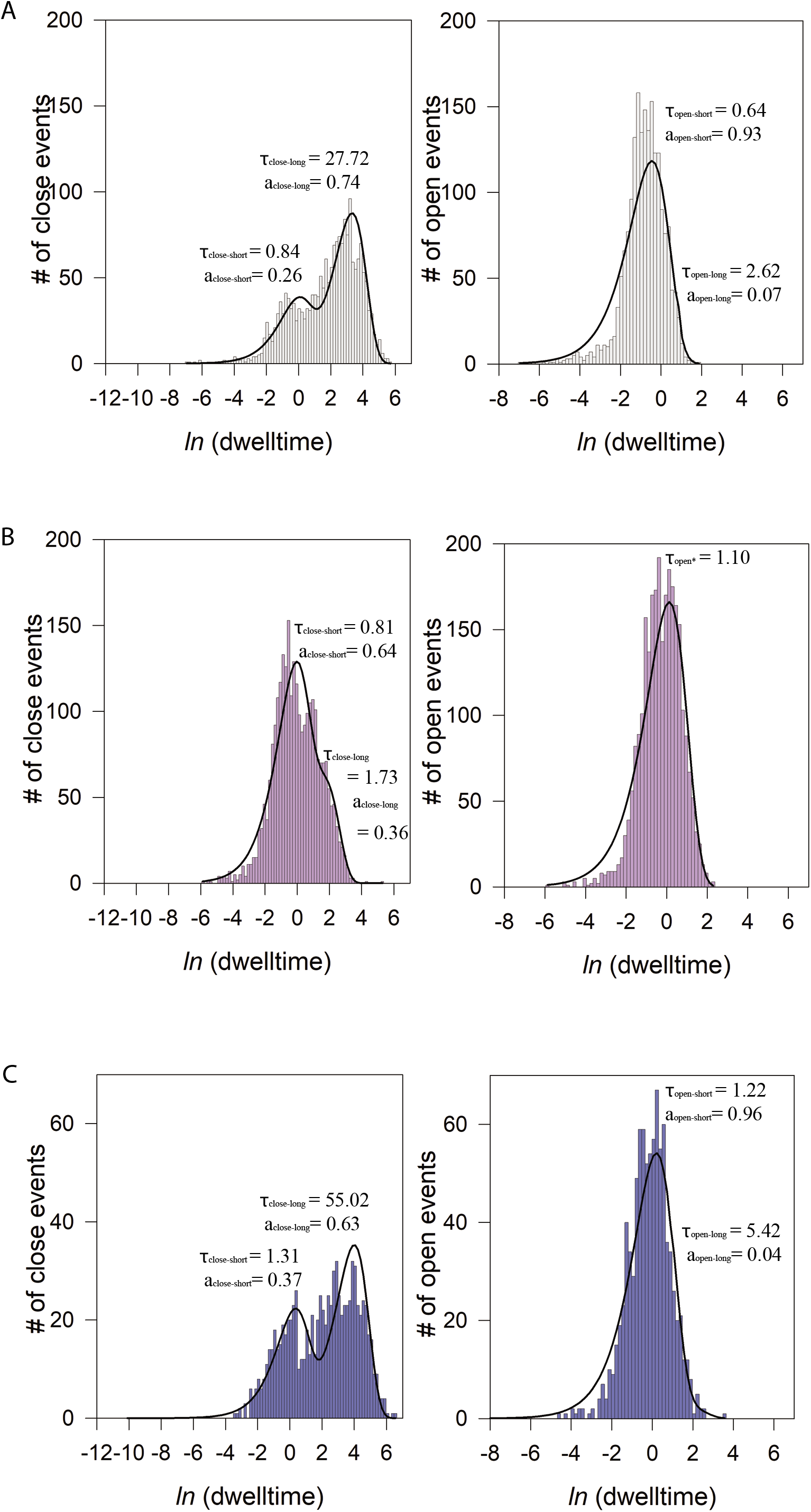
Single channel dwell time analyses. (A-C) Representative dwell time histograms from a single channel recording of TRAAK (A, gray) prior to stimulation, (B, purple) during pressure stimulation, and (C, blue) during ultrasound stimulation. Closed time histograms are shown on the left and open time histograms on the right. Maximum-likelihood fits to a four-state model with two closed and two open states are shown as black lines with mean dwell times (τ) and relative proportion (a) of total events shown for each component (closed-short, closed-long, open-short, open-long).

## METHODS

### Expression in and recording from *Xenopus laevis* oocytes

A construct encoding full length *Homo sapiens* TRAAK (UniProt Q9NYG8-1) was codon optimized for eukaryotic expression, synthesized (Genewiz), and cloned into a modified pGEMHE vector using Xho1 and EcoR1 restriction sites such that the transcribed message encodes *H. sapiens* TRAAK amino acids 1-393 with an additional amino acid sequence of “SNS” at the C-terminus. A construct encoding *Mus musculus* TASK2 (UniProt Q9JK62-1) was codon optimized for eukaryotic expression, synthesized (Genewiz), and cloned into a modified pGEMHE vector using Xho1 and EcoR1 restriction sites. The region encoding the C-terminus was truncated such that the transcribed message encodes *M. musculus* TASK2 amino acids 1-335 with an additional amino acid sequence of “SNS” at the C-terminus. cDNA was transcribed from these plasmids in vitro using T7 polymerase and 0.1–10 ng cRNA was injected into *Xenopus laevis* oocytes extracted from anaesthetized frogs. Currents were recorded at 25°C from inside-out patches excised from oocytes 1–5 days after RNA injection. Pipette solution contained (in mM) 14 KCl, 2 MgCl_2_, 10 HEPES, pH = 7.4 with KOH. The solution in the bath and US chamber contained 140 KCl, 2 MgCl_2_, 10 HEPES, 1 EGTA, pH = 7.1 with KOH. Currents were recorded using an Axopatch 200B Patch Clamp amplifier at a bandwidth of 1 kHz and digitized at 500 kHz. Single-channel patches were identified as long (minimum 3 min) recordings without superimposed channel openings after pressure-induced increase in open probability. To evaluate channel open and close durations, currents from patches containing no superimposed channel openings were unfiltered in order to preserve very brief openings that are characteristic of TRAAK. Singles were idealized by half-amplitude threshold crossing. All single channel data was analyzed using custom written software^46^.

### TRAAK reconstitution and recording in proteoliposomes

Mouse TRAAK (UniProt O88454-1) was cloned and expressed in *Pichia pastoris* cells as previously described^47^ with modifications described here. The construct used for purification included an additional 26 amino acid N-terminal sequence from Q9NYG8-1 that improved heterologous expression. The final construct is C-terminally truncated by 97 amino acids, incorporates two mutations to remove N-linked glycosylation sites (N81Q/N84Q), and is expressed as a C-terminal PreScission protease-cleavable EGFP-10xHis fusion protein. As a result, there is an additional amino acid sequence of “SNSLEVLFQ” at the C-terminus of the final purified protein after protease cleavage.

Frozen Pichia cells expressing TRAAK were disrupted by milling (Retsch model MM301) 5 times for 3 min at 25 Hz. All subsequent purification steps were carried out at 4 °C. Milled cells were resuspended in buffer A (in mM) 50 Tris pH 8.0, 150 KCl, 1 EDTA 0.1 mg/ml DNase1, 1 mg/ml pepstatin, 1 mg/ml leupeptin, 1 mg/ml aprotinin, 10 mg/ml soy trypsin inhibitor, 1 mM benzamidine, 100 μM AEBSF, 1 μM E-64, and 1 mM phenylmethysulfonyl fluoride added immediately before use) at a ratio of 1 g cell pellet per 4 ml lysis buffer and sonicated for 4 minutes with a 25% duty cycle. The solution was ultracentrifuged at 150,000 xg for 1hr at 4 °C. Pellets were transferred to a Dounce homogenizer in buffer B (buffer A + 1%DDM/0.2%CHS). Detergent was added from a 10%DDM/2%CHS stock in 200 mM Tris pH 8 that was sonicated until clear. Following homogenization, solutions were stirred for 2h at 4 °C followed by centrifugation at 35,000g for 45 min. Anti-GFP nanobody resin washed in Buffer B was added to the supernatant at a ratio of 1 mL resin = 1 mg purified anti-GFP nanobody / 15 g *Pichia* cells and stirred gently for 2 h. Resin was collected on a column and serially washed in Buffer C (buffer A + 0.1%DDM/0.02%CHS), Buffer D (buffer A + 150 mM KCl + 0.1%DDM/0.02%CHS). The resin was resuspended in 2 volumes of Buffer C with 1 mg purified Precission protease and gently rocked in column overnight. Cleaved TRAAK was eluted in ~4 column volumes of Buffer C, concentrated (50 kDa MWCO), and applied to a Superdex 200 column (GE Healthcare) equilibrated in Buffer E (20 mM Tris pH 8.0, 150 mM KCl, 1 mM EDTA, 0.025%/0.005% DDM/CHS). Peak fractions were pooled and concentrated to ~1 mg/mL for reconstitution.

Purified TRAAK was reconstituted in L-α-phosphatidylcholine extract from soybean lipids as described^48^. Proteoliposomes were snap-frozen in liquid nitrogen and stored at −80°C in De/Rehydration (DR) buffer composed of 200 KCl, 5 HEPES-KOH pH to 7.2. When preparing proteoliposomes for patching, samples were thawed at room temperature and dried 2.5-3 hours in a vacuum chamber to dehydrate. The dehydrated proteoliposomes were then rehydrated with DR buffer. Currents were recorded at 25°C from inside-out patches excised from proteoliposomes at least 12 hours after rehydration. Pipette solution contained: 5 HEPES, 20 KCl, 180 NaCl, pH 7.2 adjusted with NaOH. Bath solution contained: 5 HEPES, 200 KCl, 40 MgCl_2_, pH 7.2 adjusted with KOH. Currents were recorded using an Axopatch 200B Patch Clamp amplifier at a bandwidth of 1 kHz and digitized at 500 kHz.

### Ultrasound setup and application from inside patches

Inside out patches excised from either oocytes or proteoliposomes were quickly (within 5-10 seconds) transferred to the ultrasound chamber. The patch was centrally positioned approximately 1 inch over the transducer, separated only by bath solution. An ultrasound wave was generated using a focused immersion ultrasonic transducer, V326-SU (Olympus), which had a focus at 25.222 mm (0.993 in), nominal element size of 9.522 mm (.375 in), and an output center frequency of 4.78 MHz. A function generator (Agilent Technologies, model 33220A) was used to trigger the transducer’s ultrasound pulses. A RF amplifier (ENI, model 403LA), which receives an input voltage waveform from the function generator, provides the output power to the ultrasound transducer for producing the acoustic pressure profile of a stimulus waveform. The timing of the ultrasound stimuli was controlled by triggering the functional generator manually or by software (Clampex 10.7). In the case of ultrasound pulse generation though software, a Clampex 10.7 generated waveform triggered a first functional generator through a digitizer (Axon Digidata 1550B), which triggered a second functional generator, which triggered the RF amplifier that drives the ultrasound transducer. Solutions were degassed to minimize microbubble cavitation and ultrasound attenuation.

### In utero electroporation

Electroporation was performed on pregnant CD1 (ICR) mice (E15, Charles River ca. SC:022) as described^49^. For each surgery, the mouse was initially anesthetized with 5% isoflurane and maintained with 2.5% isoflurane. The surgery was conducted on a heating pad, and warm sterile PBS was intermittently perfused over the pups throughout the procedure. A micropipette was used to inject ~1 μl of recombinant DNA at a concentration of 2 μg/μl and into the left ventricle of each embryo’s brain (typically DNA encoding TRAAK was doped with plasmid expressing GFP at a concentration of 1:3 to facilitate screening for expression after birth). Fast-green (Sigma-Aldrich) was used to visualize a successful injection. Following successful injection, platinum plated forceps-type electrodes (5mm Tweezertrodes -BTX Harvard Apparatus) connected to the negative pole were used to gently grab both sides of the embryo’s head and the third electrode connected to the positive pole was placed slightly below lambda^50^. An Electro Square Porator (BTX Harvard Apparatus) was used to administer a train of 6 × 40 mV pulses with a 1 s delay. After the procedure, the mouse was allowed to recover and come to term, and the delivered pups were allowed to develop normally. On the day of birth, animals were screened for location and strength of electroporation by trans-cranial epifluorescence under an Olympus MVX10 fluorescence stereoscope.

### Slice electrophysiology

We used radial slices from the somatosensory barrel cortex cut along the thalamocortical plane or coronal cortical sections. The hemisphere was trimmed on both the anterior and posterior side of barrel cortex with coronal cuts, placed on its anterior side and a cut was made with a scalpel so that much of barrel cortex lay in a plane parallel to cut. The surface of this last cut was glued to the slicer tray. The preparation was aided by the use of epifluorescent goggles to visualize the expressing area. Six 300 μm slices were prepared. Cortical slices (400 μm thick) were prepared as described^51^ from the transfected hemispheres of both male and female mice aged P15-P40 using a DSK Microslicer in a reduced sodium solution containing (in mM) NaCl 83, KCl 2.5, MgSO4 3.3, NaH2PO4 1, glucose 22, sucrose 72, CaCl2 0.5, and stored submerged at 34 °C for 30 min, then at room temperature for 1–4 h in the same solution before being transferred to a submerged recording chamber maintained at 25°C in a solution containing (in mM) NaCl 119, KCl 2.5, MgSO4 1.3, NaH2PO4 1.3, glucose 20, NaHCO3 26, CaCl2 2.5. Pipettes were filled with potassium gluconate based internal solution (K-gluconate 135, NaCl 8, HEPES 10, Na3GTP 0.3, MgATP 4, EGTA 0.3). Currents were recorded using an Axopatch 200B Patch Clamp amplifier at a bandwidth of 1 kHz and digitized at 500 kHz. The recording chamber contained a cortical slice resting over a thin film of mylar. Directly under the mylar was a 1-inch tube leading to the surface plane of the ultrasound transducer.

### Ultrasound induced temperature changes

Temperature changes generated by stimulation with the 5 MHz immersion focus ultrasonic transducer (V326-SU, Olympus) were measured over time with an immersed thermocouple at the position of maximum ultrasound power. The functional generator was set to 1.0 V, 5 MHz sine wave, and infinite cycles. These measurements corresponded well to expected temperature increases calculated with the relationships{Nyborg:1998gq}:

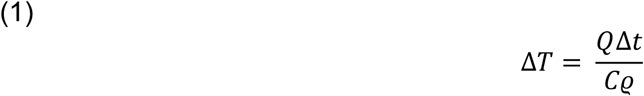

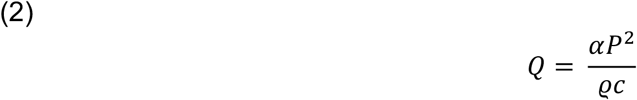

Where *ΔT* is temperature change, *Q* is ultrasound-generated heat, *Δt* is time of ultrasound stimulation, *C* is solution specific heat capacity (3600 J kg^−1^ K^−1^), *ϱ* is solution density (1028 kg m^−3^), *α* is ultrasound absorption coefficient (20 m^−1^), *P* is effective ultrasound pressure (calculated as 0.707 times peak amplitude of the sine wave), and c is the speed of sound in solution (1515 m s^−1^). These relationships were subsequently used to estimate temperature increases and design protocols that minimized heating.

### Calculating ultrasound pressure and power intensity

The output pressures were measured using a calibrated hydrophone (Onda, model HNR-0500). The hydrophone measurements were performed at the position of peak spatial pressure. When converting the measured voltages into pressures, we accounted for the hydrophone capacitance according to the manufacturer’s calibration. Using the appropriate conversion factor listed under the ‘Pa/V’ (Pascals per volt) column on the look-up table that was supplied with the calibrated hydrophone, the hydrophone voltage trace waveform was transformed into an acoustic pressure waveform measured in MPa.

We calculated the ultrasound power intensity in W/cm^2^ with the following equation:

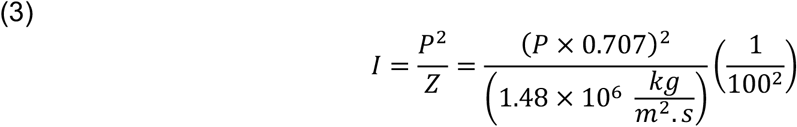

Where *P* is effective ultrasound pressure (calculated as 0.707 times peak amplitude of the pressure wave) and *Z* is the acoustic impedance (1.48×10^6^ kg m^−2^ s^−1^ was used).

### Animals

Animal procedures were reviewed and approved by the Animal Care and Use Committee at the University of California, Berkeley (AUP 2016-09-9174, AUP 2014010-6832, and AUP-2015-04-7522-1).

## Acknowledgements

We thank Dr. L. Csanády for providing software for single channel analyses; Dr. E. Iscaoff, Dr. Y. Yu, and C. Stanley for providing *Xenopus* oocytes; Dr. M. Gajowa and Dr. S. Sridharan for advice on slice recordings; I. Tayler for assistance with molecular biology; and members of the Brohawn lab for critical review and discussions. SGB and HA are New York Stem Cell Foundation-Robertson Neuroscience Investigators. This work is supported by the New York Stem Cell Foundation (SGB and HA), NIGMS grant DP2GM123496-01 (SGB), a McKnight Foundation Scholar Award (SGB), a Klingenstein-Simons Foundation Fellowship Award (SGB), and a Rose-Hill Innovator Award (SGB and HA).

## Author contributions

Conceptualization, S.G.B., H.A., and B.S.; Methodology, S.G.B., H.A., and B.S.; Investigation, BS, RAR, and K.G.; Writing – Original Draft, B.S. and S.G.B.; Writing – Review & Editing, B.S., R. A.R., K.G., H.A. and S.G.B.; Funding Acquisition, H.A. and S.G.B.; Supervision, H.A. and S. G.B.

## Notes

### Competing Interest Statement

The authors have declared no competing interest.

